# RNA-seq of seven non-neoplastic/inflammatory lymphadenitis entities identify potential molecular diagnostic markers

**DOI:** 10.1101/2025.05.29.655793

**Authors:** Julia Wang, Brian Lockhart, Jason Xu, Vinodh Pillai

## Abstract

Lymphadenopathy is a common mode of presentation for various disorders. While malignancy must be considered, there is a broad spectrum of causes that are infectious or of autoimmune origin. Clinical testing and morphological assessment are required to distinguish between neoplasms and benign pathologies. However, many causes of lymphadenopathy have similar histological presentations, making definitive diagnoses and categorization challenging. While DNA sequencing have enabled the accurate diagnosis of certain disorders, such as the genetic origin of ALPS (autoimmune lymphoproliferative syndrome), other causes of lymphadenopathy such as benign sinus histiocytosis, or Rosai-Dorfman disease (RDD), cannot be diagnosed with current technologies. Likewise, several modes of infection, including *Toxoplasmosis gondii*, Epstein-Barr virus (EBV), and *Bartonella henselae* (Cat-scratch disease) require further investigation. RNA sequencing enables the assessment of differentially expressed genes (DEG) that reveal up or downregulated pathways. These DEG signatures and pathways can help differentiate lymphadenopathies that appear similar histologically. In this study, RNA sequencing of 46 cases of lymphadenopathies were performed and analyzed. ALPS and RDD cases provided confirmation that RNA sequencing can identified expected gene expression and activated pathways. Our results showed the unique activation of the interferon pathway was identified in sinus histiocytosis cases. Through advances in RNA sequencing, potential genetic signatures can be discovered in a gamut of varying conditions.

## Introduction

Lymphadenopathy can arise from a variety of origins: infection, autoimmune, or malignancy. The diagnostic work-up for many of these causes is similar -- a morphological assessment and several laboratory tests are conducted. Reactive or benign lymphadenopathy are especially challenging to diagnose and treat in persistent cases due to lack of understanding of the pathogenesis.

Many genetic markers of malignancy have been heavily characterized due to their severity^1^, but genetic signatures of certain infectious and benign lymphoproliferations are unclear. While it is possible to confirm an infection by a simple blood test, the factors leading to nodal lymphoid response is not completely understood. An additional challenge arises with the limited tissues obtained during the increasingly utilized core biopsies. The overall architecture and focal findings are critical for morphological diagnosis of several reactive lymphadenopathies. If each entity had a unique set of molecular findings, there may be an improvement in diagnostic accuracy.

Mononucleosis infection caused by the Epstein Barr Virus (EBV) has been found to elicit response by CD8+ lymphocytes^2^. Conversely, the mechanism of immune response during infection by *Bartonella henselae* (Cat-scratch disease) and *Toxoplasmosis gondii* are not well described. There are many descriptions of the morphological changes in these conditions, but contributory genetic markers have yet to be described. This also applies to benign lymphoproliferations such as sinus histiocytosis and Rosai-Dorfman disease (sinus histiocytosis with massive lymphadenopathy). An established lymphoproliferative disorder, Autoimmune lymphoproliferative syndrome (ALPS), is characterized by the overproduction of T lymphocytes that are CD4, CD8 negative^3^. This is contributed to a deletion of the *Fas* gene, which prevents these cells from undergoing apoptosis^3^.

To better understand the genetic signatures of these disorders, a cohort of inflammatory lymph nodes were selected from the Children’s Hospital of Philadelphia pathology archives and formalin fixed paraffin tissue (FFPE) was obtained for RNA sequencing. With a more thorough understanding of lymphadenopathy gene signatures, diagnosis may be possible with core biopsies and resection may be avoided.

## Methods

### Cases and controls

All cases (n=46) were identified through the Children’s Hospital of Philadelphia pathology archives. Specimens were separated into two cohorts, “BL002” and “BL003”. BL002 consisted of ALPS, RDD, and sinus histiocytosis (SH) samples, while BL003 consisted of EBV, *Toxoplasmosis gondii*, Cat-scratch lymphadenitis, and dermatopathic effect samples. To confirm diagnoses, pertinent laboratory testing, as well as thorough pathologic evaluation, were pursued prior to case selection. All cases were derived from lymph nodes, and formalin fixed paraffin embedded (FFPE) blocks were cut to generate FFPE tissue slides. The date range for these specimens spanned from 1994 to 2021. A positive control was utilized in experiment BL002, while dermatopathic effect cases served as an internal control for analysis.

### RNA sequencing

Targeted RNA sequencing analysis was performed on archived FFPE tissues using the HTG EdgeSeq platform (HTG Molecular Diagnostics, Arizona) as per the instruction manual. Compared to other RNA sequencing technologies, the HTG system has a high success rate with FFPE tissue^4^. FFPE tissue was scraped from slides, and correctly prepped for quantitative nuclease protection on the HTG Edgeseq Processor. A panel consisting of 2003 immune response genes (EdgeSeq Immune Response Panel) was selected for the expression analysis of these samples. Select samples from both BL002 and BL003 (ALPS and *Toxoplasmosis* cases, respectively) were also sequenced via the whole transcriptome panel. Following processing, samples were amplified via PCR (ProFlex PCR system, Applied Biosystems) and Illumina primer tags were added. The sample libraries were cleaned and quantified using qPCR (BioRad CFX Connect Real Time System). Libraries were then loaded onto a NextSeq 550/500 (single end, 75bp, read length) for RNA sequencing (Illumina USA). Data was extracted and parsed by the HTG specific parser. Parsed data underwent differential expression (DE) by HTG DeSeq2, a python-based platform specific to HTG Molecular Diagnostics, Inc. Gene expression pathway analysis was completed by the online genetic database, Metascape.

### Pathway analysis

For each entity, the gene expression were compared to all other samples or dermatopathic effect samples. Genes for pathway analysis were then selected from the differential expression list based off of adjusted p-value (see results section for more details). Metascape was the online application utilized for this interpretation, which attributes the genes to certain pathways, cell lines, and genetically related clusters^5^. The background gene list submitted to Metascape for analysis is defined as the 2000 genes panel (HTG EdgeSeq Immune Response Panel).

## Results

### Case characteristics

Basic demographic of the 46 cases selected are listed in Table 1. Of the six ALPS cases, follicular hyperplasia was noted as a histological feature (Figure 1a) in all cases except for in case ALPS4, where the patient also exhibited features of Rosai-Dorfman Disease. All six cases had at least one positive ALPS flow cytometric screen where the percentage of double negative T cells out of all T cells was greater than 2.5% or the absolute number was above 45 cells/uL. All cases had significant cytopenias and two cases had splenomegaly and subsequent splenectomy. Four cases had a concurrent diagnosis of Evans syndrome. Two cases had a diagnosis of Common Variable Immune Deficiency. Three cases had genetic testing (Autoimmune Lymphoproliferative Syndrome Gene Sequencing Panel) done at Cincinnati Children’s Hospital and all were negative. One case was found to have decreased Fas-mediated apoptosis on functional test (also done at Cincinnati Children’s Hospital).

**Table 1:**
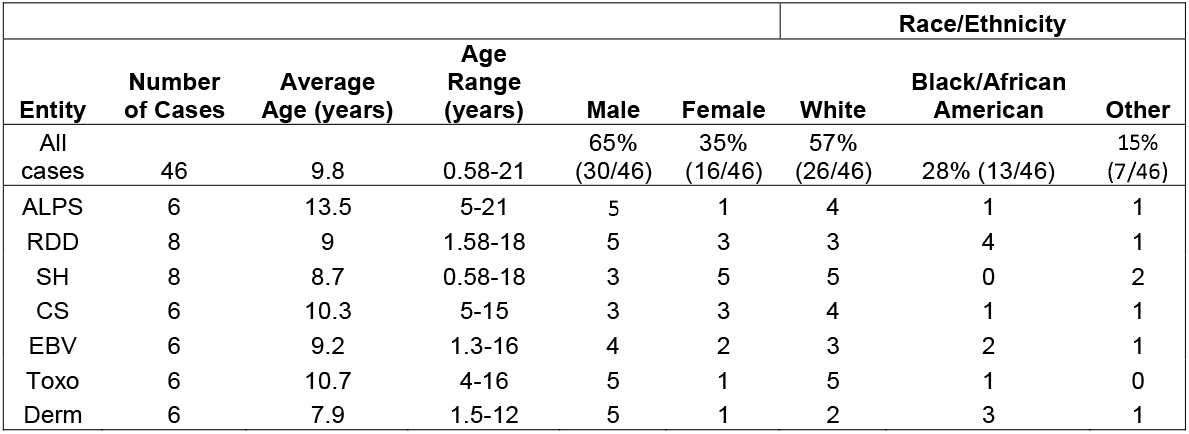

**Figure 1a:**
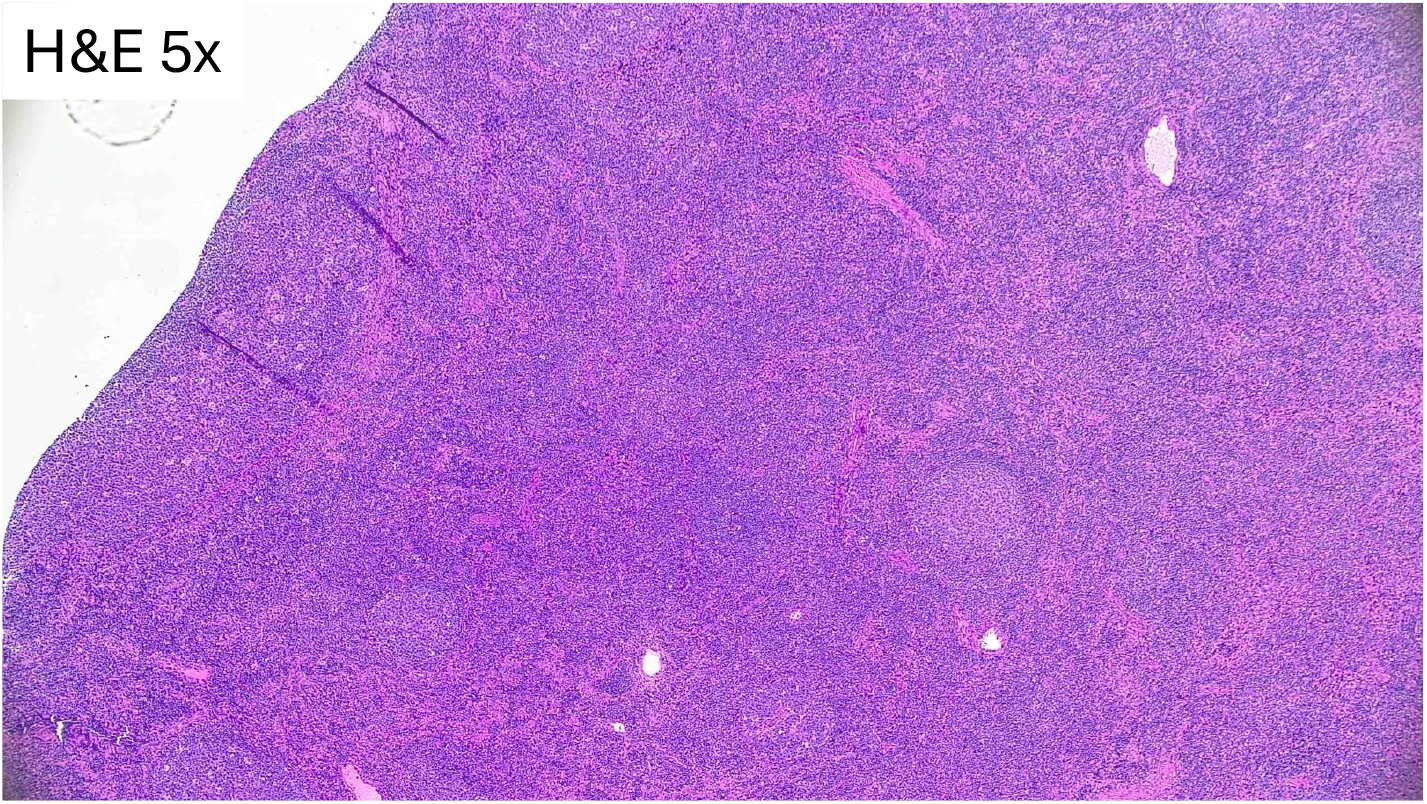
Histology (H&E) for ALPS

Of the eight cases of RDD, four cases had next generation sequencing performed on the lymph node with one case showing low level *BRAF* p.V600E mutation and the others were negative. On histology, emperipolesis was present in call cases (Figure 1b). Eight cases of sinus histiocytosis (Figure 1c) are included in this study, notably, two cases have a history of RDD that was not present on the specimen included in this study. Three out of eight cases had concurrent diagnosis of an autoimmune-related disease. Of the six *Bartonella henselae* (Cat-scratch disease) cases (Figure 1d), five cases had positive IgG serology with a range of 1:128 to 1:1024. Two cases had confirmed contact with cats. Of the six cases of EBV (mononucleosis) (Figure 1e), EBER (ISH) was positive in all cases and atypical lymphocytes were noted in the peripheral blood of three cases. In all six cases of toxoplasmosis (Figure 1f), serology testing were performed and was positive in one case. No documentation of exposure to cats were noted in any of the cases. Classic histology triad was present in five out of six cases. Six cases of dermatopathic change were included to serve as the control group (Figure 1g).

**Figure 1b:**
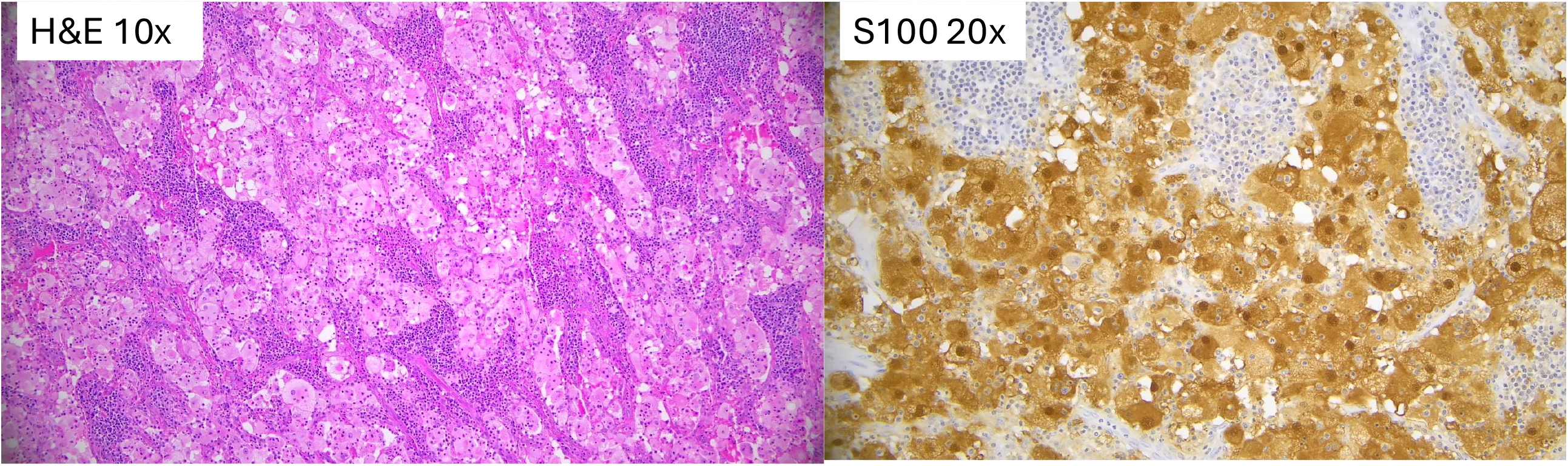
Histology for RDD

**Figure 1c:**
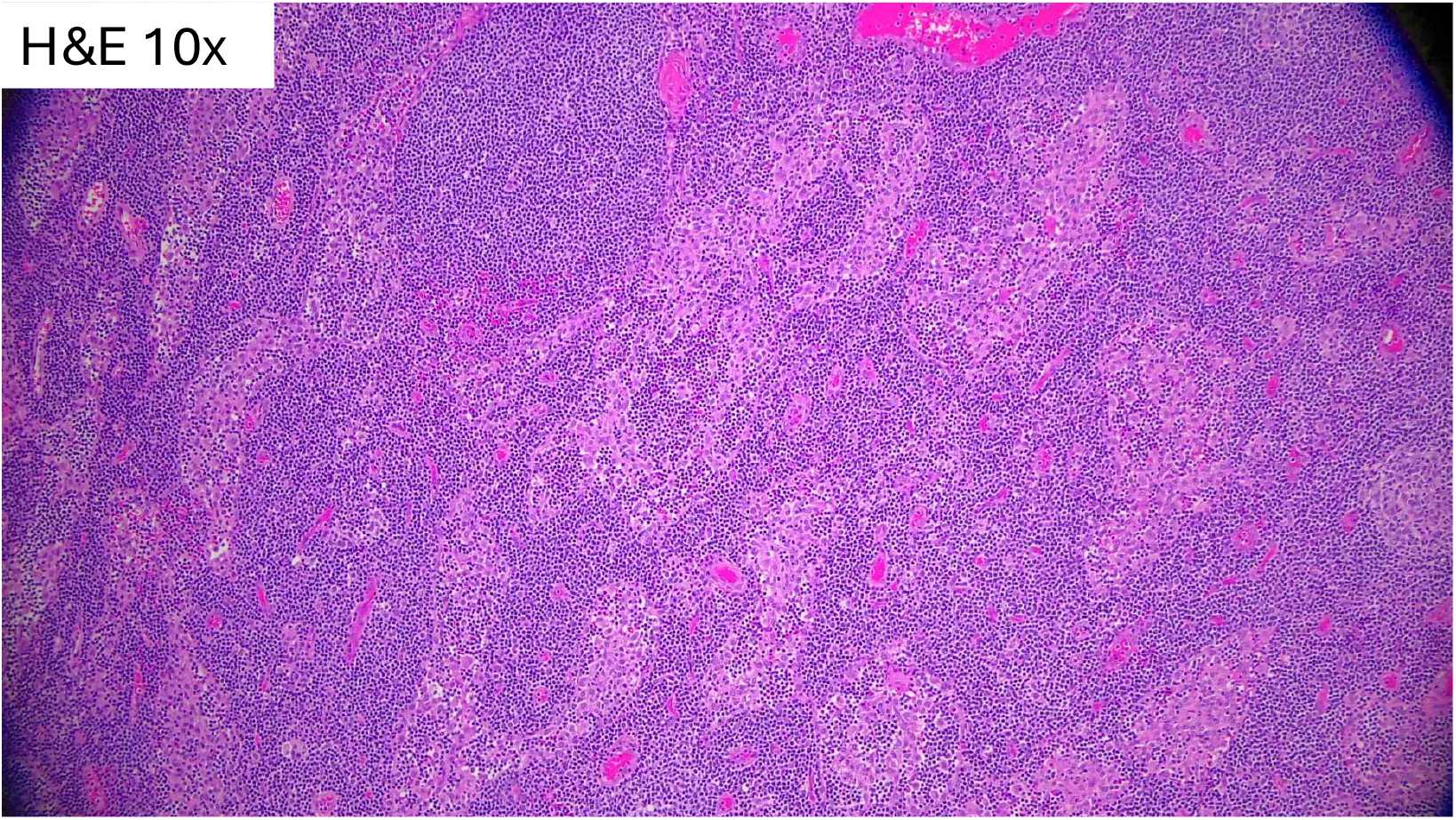
Histology (H&E)for SH (Sinus Histiocytosis)

**Figure 1d:**
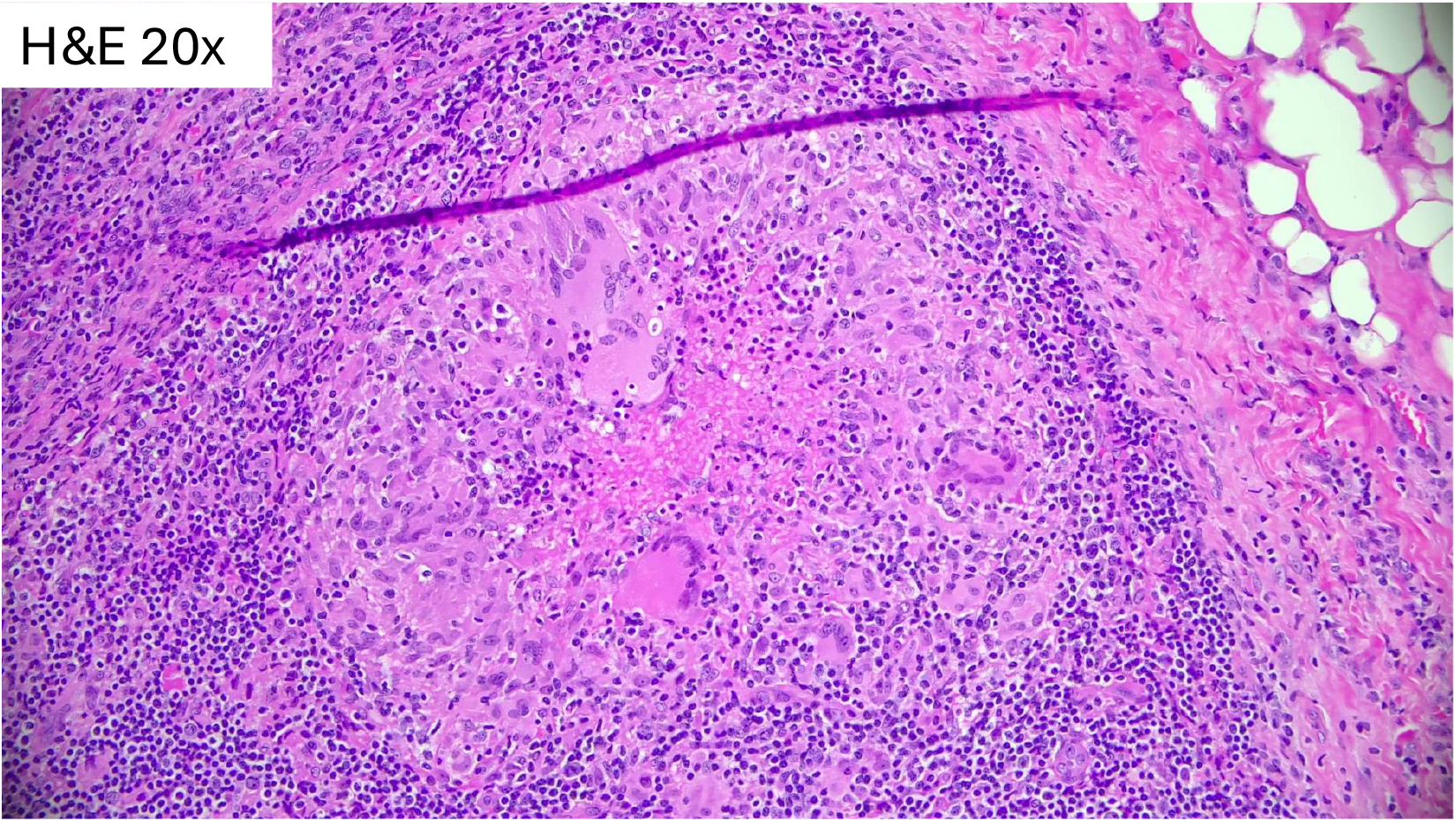
Histology (H&E)for CS (Cat-scratch disease)

**Figure 1e:**
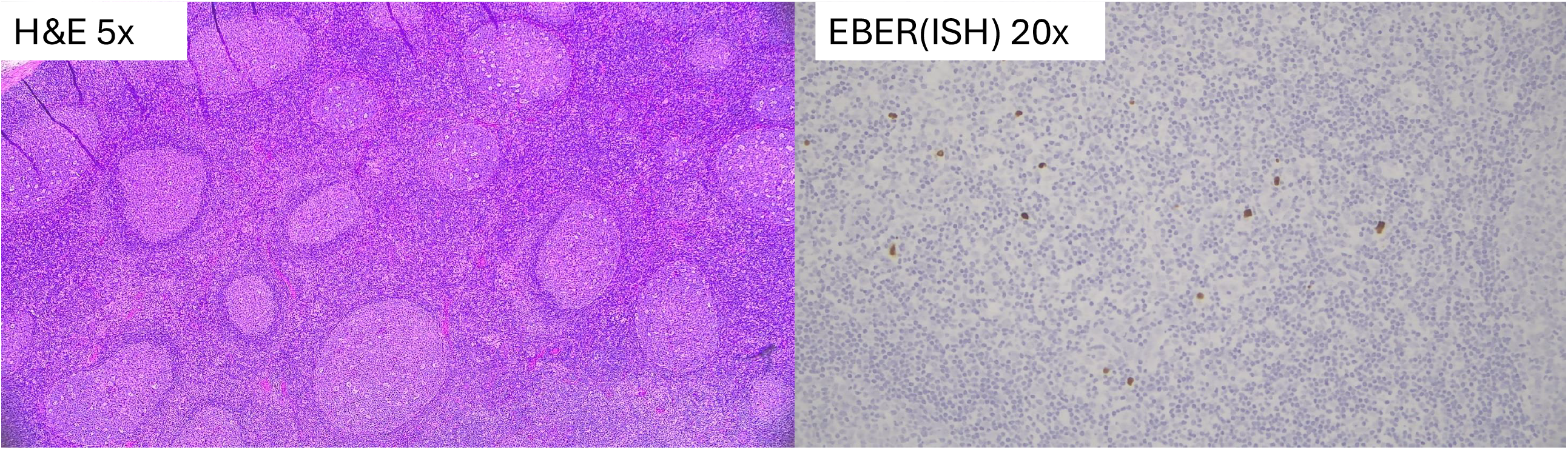
Histology (H&E)for EBV lymphadenitis

**Figure 1f:**
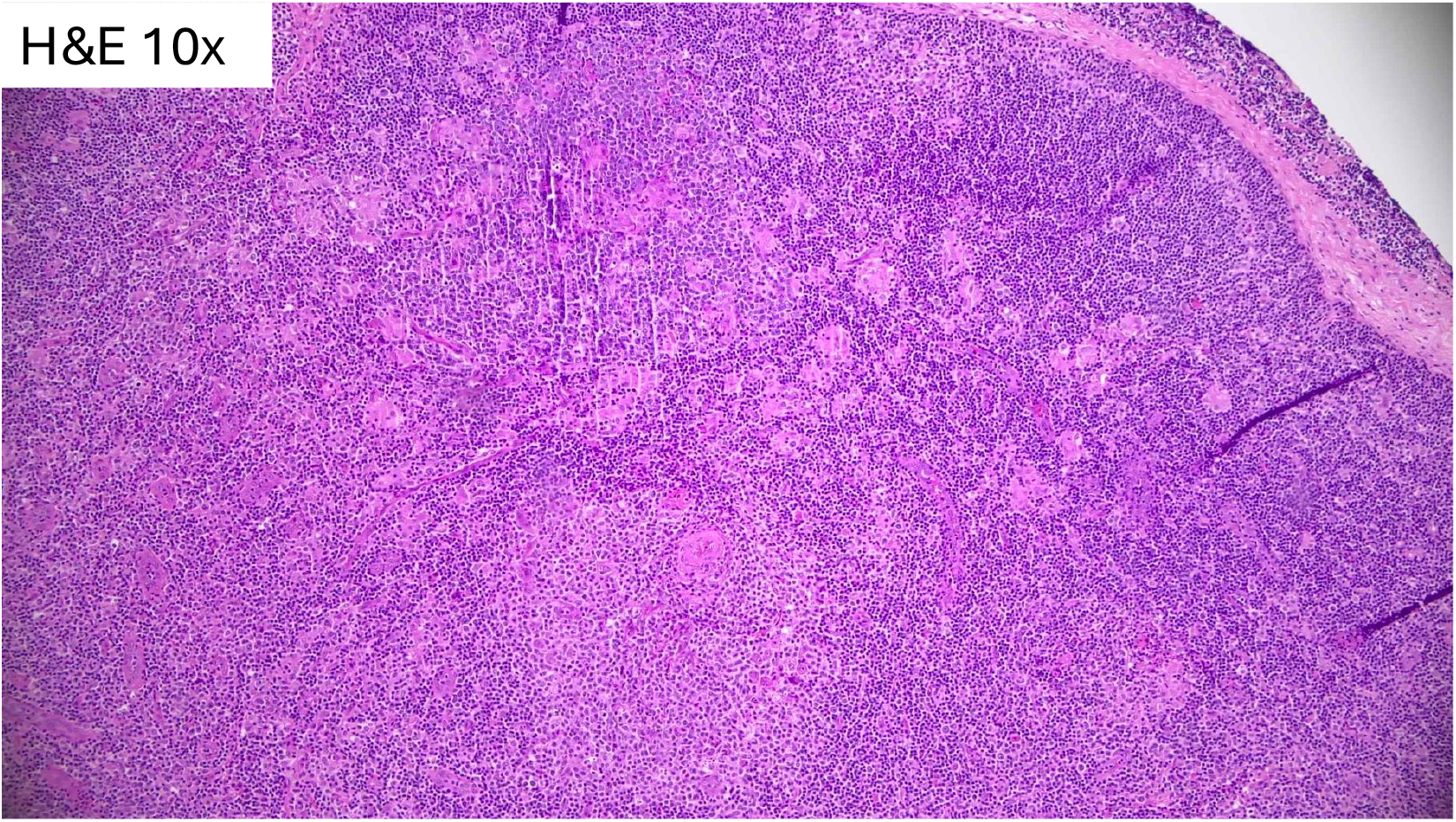
Histology (H&E)for Toxoplasmosis lymphadenitis

**Figure 1g:**
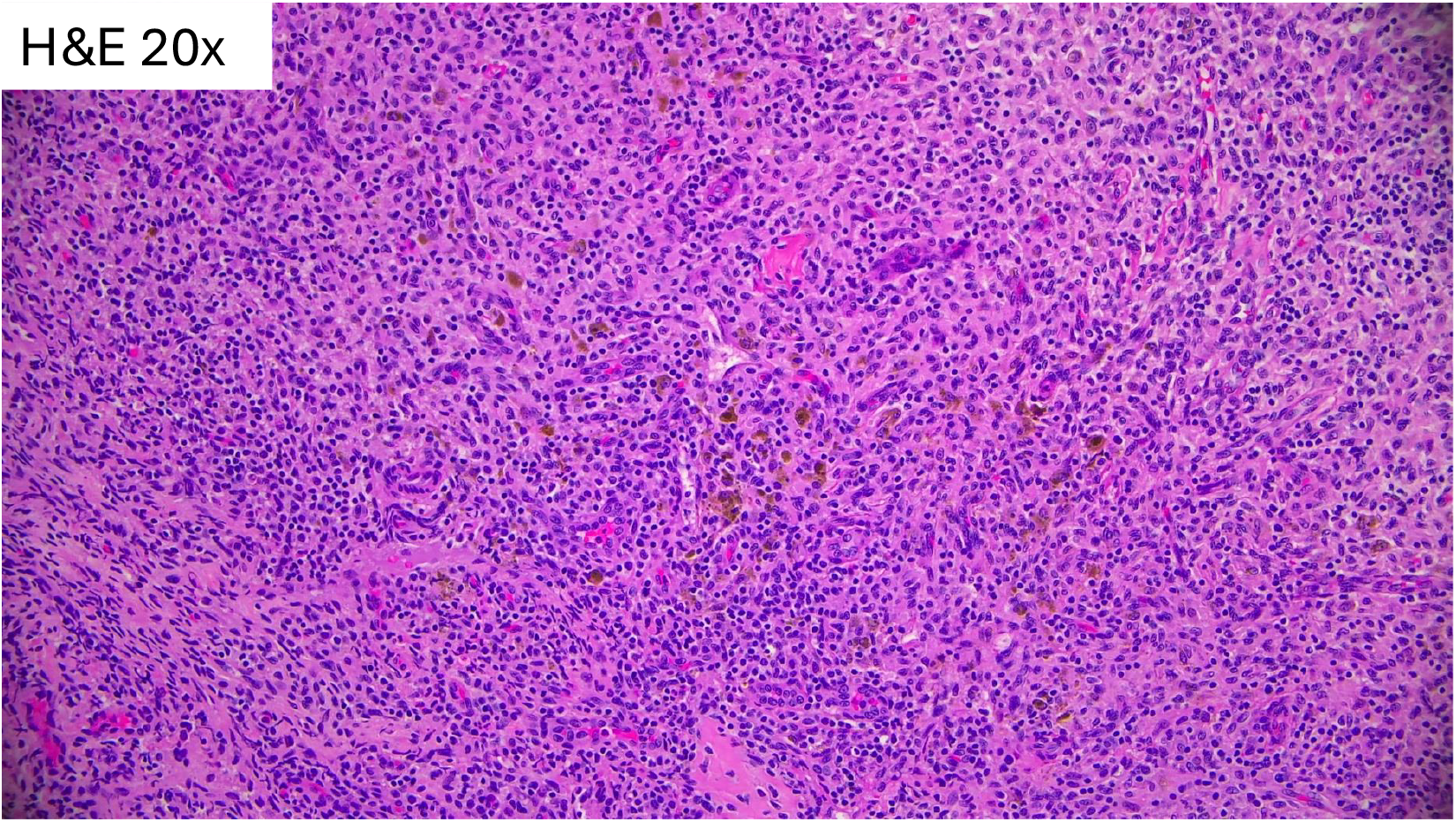
Histology (H&E)for Dermatopathic effect

The DEGs in RDD cases were compared with all other samples (Figure 2a). In RDD, 72 genes were significantly down-regulated (adjusted p-value < 0.1) and 51 genes were significantly up-regulated when compared to all other groups (see Supplemental ***). Within the 51 genes upregulated in RDD, IL-10 signaling is among the top Gene Ontology biological process. Pathways that are significantly represented in the 72 down-regulated genes include cell cycle related pathways including chromosome organization and CKAP4 and toll-like receptor signaling pathways.

**Figure 2a:**
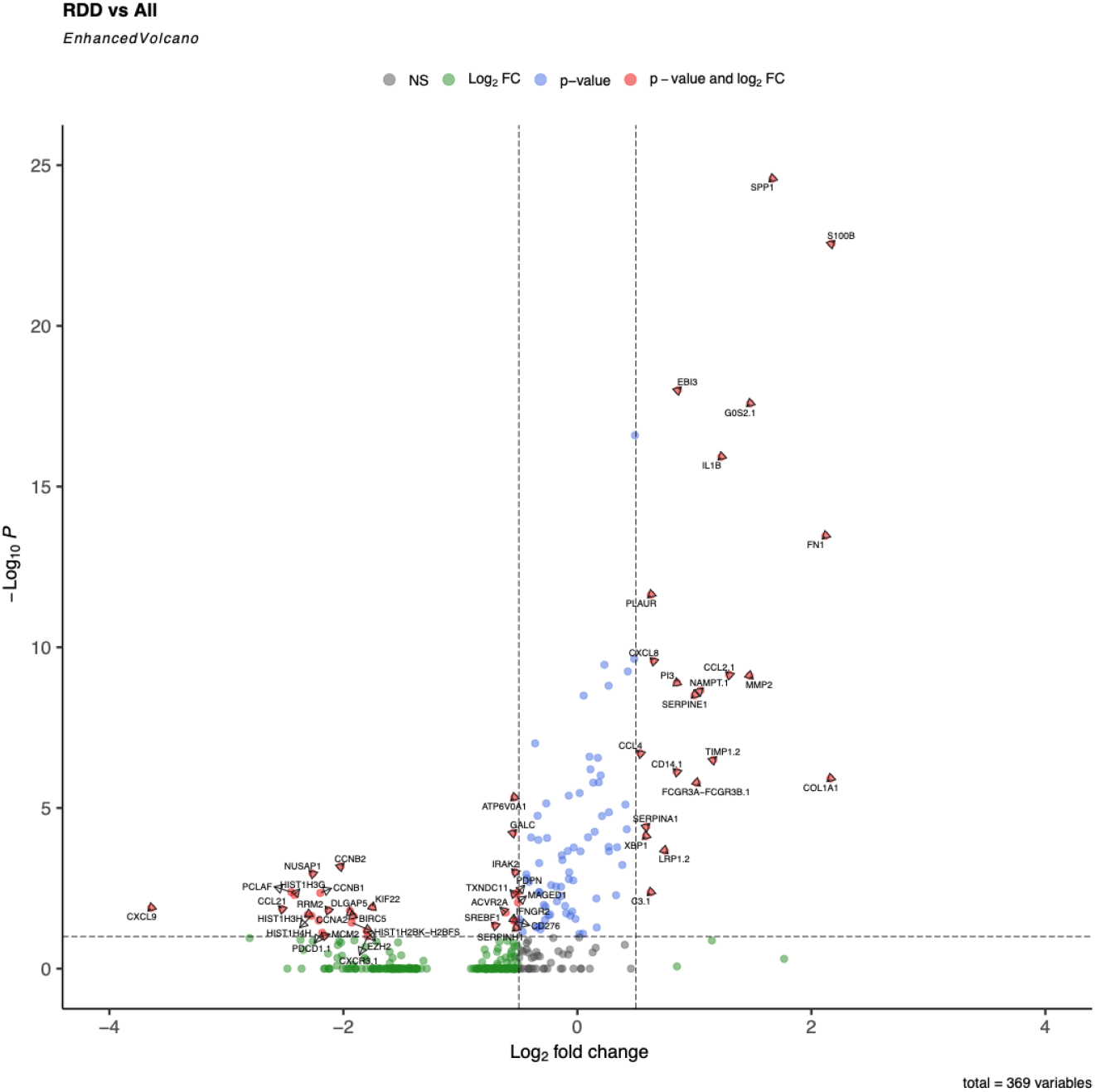
Volcano plot for RDD vs all other cases

In SH, 3 genes, *CYP27A1, ESRRA*, and *SPP1* were significantly down-regulated (adjusted p-value < 0.01) and 20 genes were significantly up-regulated when compared to all other groups (Figure 2b). The most significantly up-regulated genes in SH included *ISG15, IFI44*, and *IFI44L*. Pathway analysis revealed that interferon alpha/beta signaling and antiviral processes are the most significantly up-regulated in SH.

**Figure 2b:**
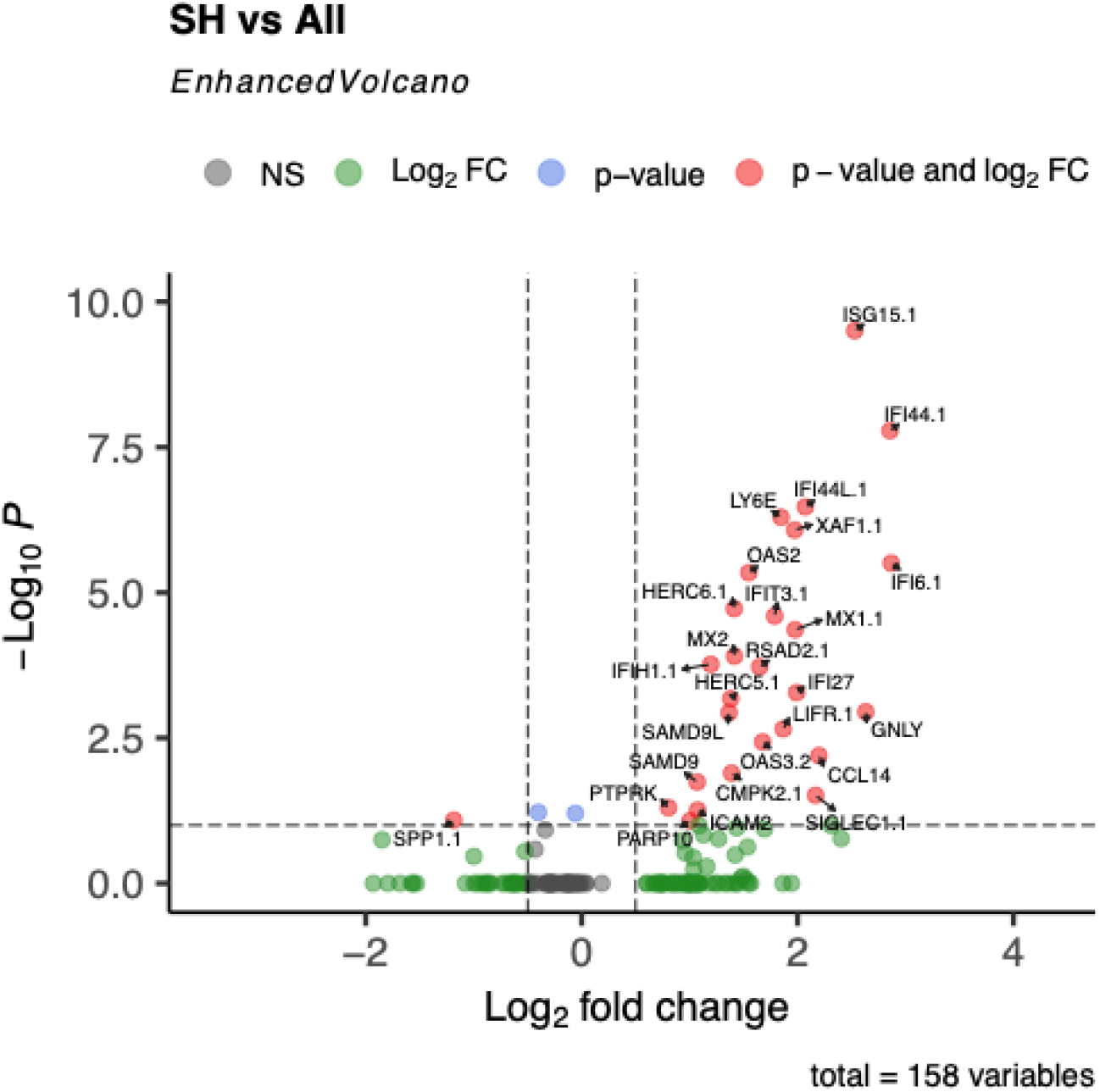
Volcano plot for SH vs all other cases

In ALPS, three genes, *PADI2, PCL-1*, and *PCL-2*, were significantly down-regulated (adjusted p-value < 0.25) when compared to all other groups.

In CS, 11 genes were significantly down-regulated (adjusted p-value < 0.25) and four genes, including *SLAMF8, IL18BP, FBP1*, and *G6PD* were significantly up-regulated when compared to all other groups (Figure 2c). Pathway analysis identified enriched ontology for “negative regulation of protein localization” and “positive regulation of peptidase activity” from the list of down-regulated genes and “cellular response to chemical stress” from the list of up-regulated genes.

**Figure 2c:**
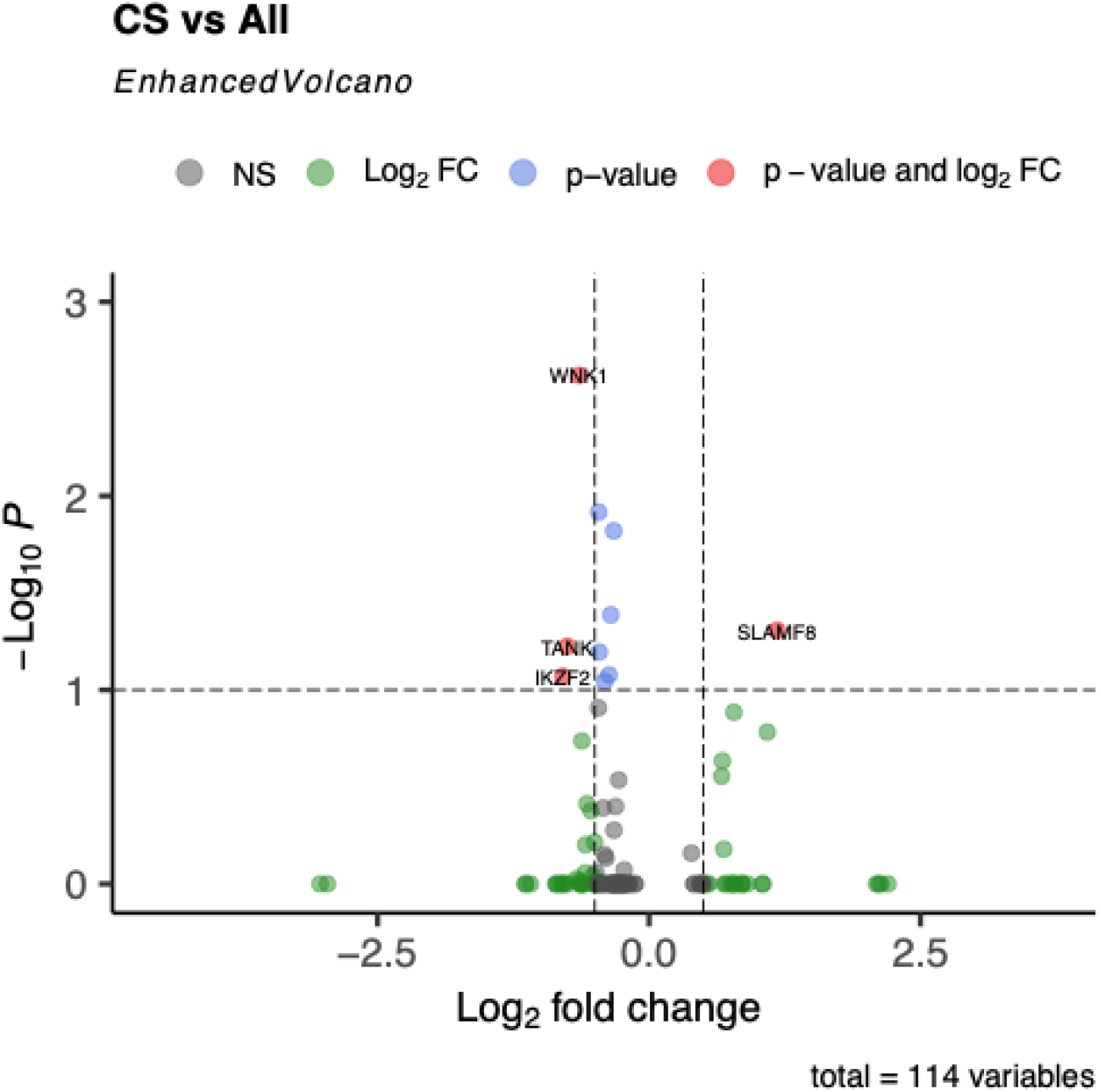
Volcano plot for CS vs all other cases

In EBV, 7 genes were significantly down-regulated (adjusted p-value < 0.25) and one gene, *CD8A*, was significantly up-regulated when compared to dermatopathic samples (Figure 2d). Pathway analysis identified enrichment of gene ontologies “Polycystic Ovary Syndrome,” “cell chemotaxis,” and “Dermatitis” from the list of significantly down-regulated genes. *CCL5* was significantly down-regulated (adjusted p-value < 0.25) when compared to all other groups.

**Figure 2d:**
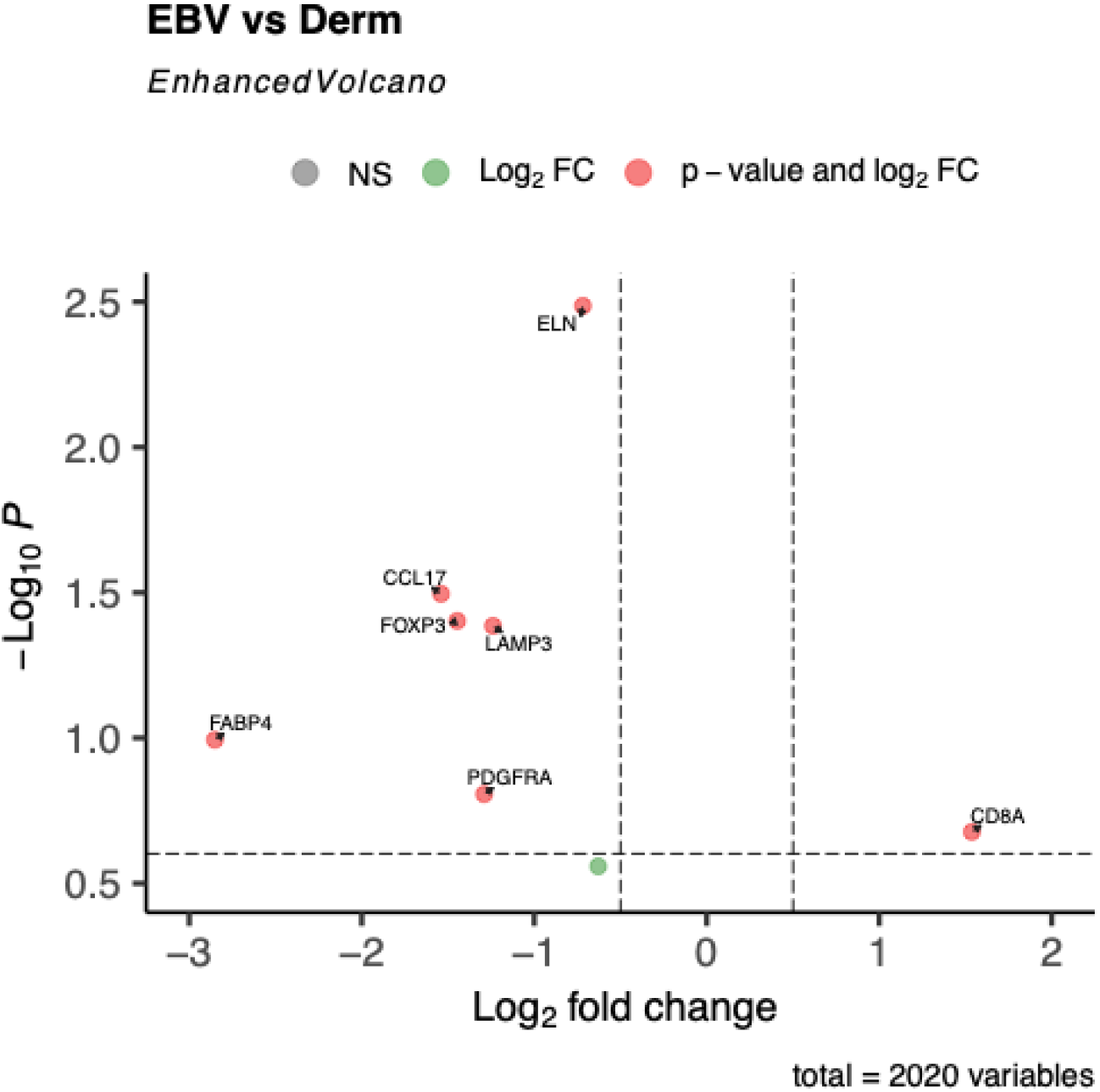
Volcano plot for EBV vs Derm

In toxoplasmosis, 2 genes (*MAP2K5* and *JAG1*) were significantly down-regulated (adjusted p-value < 0.25) and four genes, including *IL2RB, NR1H3, SH2B1*, and *KIR2DS2-KIR2DS4*, were significantly up-regulated when compared to all other groups.

In dermatopathic change samples, six genes (*CCL22, CCL17, LAMP3, CCR4, CCL13, FOXP3*) were significantly up-regulated (adjusted p-value < 0.25) when compared to all other groups. Pathway analysis identified enrichment of gene ontology “Chemokine receptors bind chemokines” from the list of significantly up-regulated genes.

## Discussion

### Rosai-Dorfman Disease (RDD)

Given that mutations in the mitogen-activated protein kinase (MAPK) pathway has been reported in RDD, it is not surprising that there is likely cross talk between the MAPK and IL-10 pathways. In one study, it was reported that in chronic intestinal inflammation, IL-10 inhibits the p38/MAPK-activated protein kinase-2 pathway which then inhibits TNF translation^6^. Notably, the most significantly up-regulated genes in RDD include *SPP1, S100B, and EBi3*. Secreted phosphoprotein 1 (SPP1) and S100 calcium binding protein B (S100B) are expressed in macrophages/histiocytes, which are hallmarks of RDD. The upregulation of *Epstein-Barr virus induced 3 (EBI3*) may suggest that there is a role in pathogen response in RDD pathogenesis.

Pathways represented in the down-regulated genes include cell cycle related pathways including chromosome organization and CKAP4 and toll-like receptor signaling pathways. Interestingly, this list of genes is also associated with the negative regulation of myeloid cell differentiation, which implies that by downregulating these genes in RDD, myeloid differentiation is in fact positively regulated. The CKAP4 pathway has been reported to play roles in multiple carcinomas and glioblastoma^7,8^. The most significantly down-regulated genes in RDD include *SLC11A2, HIF1A*, and *STP6V0A1. Solute carrier family 11 member 2 (SLC11A2)* is poorly studied in the immune system but is associated with iron homeostasis^9^ and gynecological cancers^10^. *Hypoxia inducible factor 1 subunit alpha (HIF1A*) is expressed in neutrophils and eosinophils and is known primarily for its role in response to hypoxia but also has significance in host-virus interaction and tumor angiogenesis^11^. *ATPase H+ transporting V0 subunit a1 (ATP6V0A1*) is strongly expressed in neuronal tissue and have low expression in immune cells^11^.

### Sinus Histiocytosis (SH)

Sinus histiocytosis, outside of Rosai-Dorfman disease, has not been previously genetically profiled. In our study, the most significantly up-regulated genes in SH include *ISG15 ubiquitin like modifier (ISG15)* which is a well-studied interferon-induced gene. ISG15 can inhibit viral replication and modulate host damage and repair response^12^. *Interferon induced protein 44 (IFI44)* and *Interferon induced protein 44 like (IFI44L)* are also interferon-inducible genes. They are primarily known as viral response genes that help to restrict viral replication and bacterial proliferation.

In SH, 3 genes, *CYP27A1, ESRRA*, and *SPP1* are significantly down-regulated. *Secreted phosphoprotein 1 (SPP1)* or osteoponin (OPN) is expressed in macrophages and has been reported to play important roles in histiocyte migration, cell adhesion, and others such as phagocytosis^13^. It is unknown if *Cytochrome P450 family 27 subfamily A member 1* (*CYP27A1)* or *Estrogen related receptor alpha (ESRRA)* play a role in histiocyte biology and pathology.

Pathway analysis revealed that interferon alpha/beta signaling and antiviral processes are the most significantly up-regulated in SH. Interestingly, three patients with Erdheim-Chester disease were successfully treated with interferon alpha and had sustained response for 3-4.5 years^14^. This is thought to be due to interferon alpha promoting the terminal differentiation of histiocytes and dendritic cells^15^. Although currently, no viral infection has been found to be associated with SH, there has been reports of viral triggers of hemophagocytic histiocytosis^16,17^. Together, these highlighted genes and pathway suggest that a viral process may be key in the pathogenesis of SH.

### Autoimmune lymphoproliferative syndrome (ALPS)

ALPS is characterized by massive lymphadenopathy, hepatosplenomegaly, autoimmunity and the increased TCR-alpha beta, CD4-CD8-T cells. The mechanism of pathogenesis is decreased T cell and B cell apoptosis attributed to heterozygous mutations of the Fas gene and other components of the same pathway. The function of *Peptidyl Arginine Deiminase 2 (PADI2)* is the citrullination of the C-terminal domain of RNA polymerase II and is known to play a role in breast cancer^18^. In the immune system, *PADI2* is known to be expressed in peripheral blood mononuclear cells and are important in the differentiation and function of T helper (Th) cells by citrullinating GATA3 and RORγt^19^. Thus, the down-regulation of *PADI2* may enhance the differentiation of type helper T (Th2) cells in ALPS. It has been reported that Th2 cells in ALPS exhibit abnormal cytokine response when faced with various in vitro stimulation, which may account for autoimmune features of ALPS^20^.

*PCL-1 (PHF1)* and *PCL-2 (MTF2)* are members of the polycomb-like proteins (PCL) family. *PCL-1 or PHD Finger Protein 1 (PHF1)* is reported to be involved in various tumors (including breast cancer and endometrial stromal sarcomas) and regulate cell growth arrest and apoptosis^21^. *PCL-2 or Metal Response Element Binding Transcription Factor 2 (MTF2)* has been shown to be highly expressed in and drive progression of liver cancer and glioma^22^. While it is unknown what role, if any, that *PCL-1* or *PCL-2* play in the immune system, there is one study with an ALPS murine model that valproic acid, a histone deacetylase inhibitor, can decrease the lymphoproliferation^23^. This points towards a role for histone modification in ALPS pathogenesis in which polycomb-like proteins may contribute to.

### Cat-scratch disease (CS)

In the literature, the only sequencing study applied to cat scratch disease was used to detect *B. Henselae* by NGS in the peripheral blood^24^. Our study reported the very first set of unique gene expression profile of this entity. *SLAMF8* has been reported to regulate macrophage microbicidal mechanism in a mice model^25^, but has not been previously associated with cat-scratch disease until our study. Future studies may help elucidate if the up-regulation of *SLAMF8* may be the driver of the lymphadenitis histology observed in this entity.

### EBV-lymphadenitis

While EBV-associated malignancies have been extensively studied and profiled genetically, EBV-lymphadenitis has not been studied and profiled by sequencing methodologies. CD8a upregulation can be explained by the critical role of EBV-specific CD8+ T cell in the response to an EBV infection^26^.

### Toxoplasmic lymphadenitis (Toxo)

(In toxoplasmosis, 2 genes (*MAP2K5* and *JAG1*) were significantly down-regulated (adjusted p-value < 0.25) and four genes, including *IL2RB, NR1H3, SH2B1*, and *KIR2DS2-KIR2DS4*, were significantly up-regulated when compared to all other groups.)

A previous study performed spatial profiling of monocytoid B cells^27^ and found a gene expression profile that does not overlap with our findings, which could be explained by the fact that our results represents the average of all cells in specifically in toxoplasmosis lymphadenitits in contrast to monocytoid B cells across different entities. *IL2RB* is well studied and implicated in immune response while NR1H3 is associated with macrophage regulation^28^.

### Dermatopathic change (Derm)

In lymph nodes with dermatopathic changes, there is up-regulation of chemokines derived from macrophages such as *CCL22*. Dermatopathic lymphadenitis has not been previously genetically profiled in the literature, therefore, this is the first report of an elevation in chemokine gene expression in this entity.

In summary, this study reports the gene expression profile of seven different inflammatory nonneoplastic lymphadenopathies. This shows the potential of using these data for diagnosing lymphadenopathies using core biopsies.

